# Rapid high throughput SYBR green assay for identifying the malaria vectors *Anopheles arabiensis*, *Anopheles coluzzii* and *Anopheles gambiae s.s. Giles*

**DOI:** 10.1101/448910

**Authors:** Joseph Chabi, Arjen Van’t Hof, Louis K. N’dri, Alex Datsomor, Dora Okyere, Harun Njoroge, Dimitra Pipini, Melinda P. Hadi, Dziedzom K. de Souza, Takashi Suzuki, Samuel K. Dadzie, Helen P. Jamet

## Abstract

The *Anopheles gambiae sensu lato* species complex consists of a number of cryptic species with different habitats and behaviours. These morphologically indistinct species are identified by chromosome banding and molecular diagnostic techniques which are still under improvement even though the current SINE method for identification between *An. coluzzii* and *An. gambiae* works reliably. This study describes a refinement of the SINE method to increase sensitivity and high throughput method for the identification of both species and *An. arabiensis* using amplicon dissociation characteristics.

Field collected samples, laboratory reared colonies and crossed specimens of the two species were used for the design of the protocol. *An. gambiae*, *An. coluzzii*, and hybrids of the two species were provided by the insectary of Vestergaard-NMIMR Vector Labs at the Noguchi Memorial Institute for Medical Research (Ghana) and *An. arabiensis* from Kenya. Samples were first characterised using conventional SINE PCR method, and further assayed using SYBR green, an intercalating fluorescent dye.

The three species and hybrids were clearly differentiated using the melting temperature of the dissociation curves, with derivative peaks at 72 Celsius for *An. arabiensis*, 75°C for *An. gambiae* and 86°C for *An. coluzzii*. The hybrids (*An. gambiae* / *An. coluzzii*) showed both peaks. This work is the first to describe a SYBR green real time PCR method for the characterization of *An. arabiensis*, *An. gambiae* and *An. coluzzii* and was purposely designed for basic melt-curve analysis (rather than high-resolution melt-curve) to allow it to be used on a wide range of real-time PCR machines.

## INTRODUCTION

The *Anopheles gambiae* s.l. (sensu lato) complex comprises of at least seven mosquito species originally defined by polytene chromosome analysis (Coluzzi *et al*. 1979, Coluzzi *et al*. 2002), and currently identified by PCR-diagnostic assays based on specific DNA nucleotide differences in the intergenic spacer (IGS) of the ribosomal DNA (rDNA) (Scott *et al*. 1993, Townson and Onapa 1994, Fettene *et al*. 2002). Detailed analyses of the IGS region of rDNA further revealed nucleotide substitutions that differentiated between the two forms within the *Anopheles gambiae s.s* complex designated as S and M molecular forms (della Torre *et al.* 2001), and were recently named *An. gambiae* and *An. coluzzii* (Coetzee *et al*. 2013). These two species can be identified by PCR and gel electrophoresis showing the presence/absence of a diagnostic Short Interspersed Nuclear Element (SINE) on the Xchromosome and are considered to be the most efficient malaria vectors in the world and particularly in Africa (Coetzee 2004).

Hybrid forms of the *An. gambiae* and *An. coluzzii* have been identified (Fanello *et al*. 2002, Santolamazza *et al*. 2008), although information on survival rates of wild populations is still lacking. However, it has been described that the first progeny (F1) of hybrids were fully fertile (Diabate 2007), and recent laboratory-based crossing experiments have shown that the hybrids of *An. gambiae* / *An. coluzzii* can be maintained over several generations (Chabi 2016).

*Anopheles gambiae* and *An. coluzzii* are characterised by a high degree of gene flow restriction, low level of genetic differentiation, and a largely overlapping geographical and temporal distribution (della Torre *et al*. 2005). Furthermore, the two species, together with *An. arabiensis,* can live in sympatry but show different living characteristics such as insecticide resistance profile with varying resistance allele distributions (Awolola *et al*. 2005). Therefore, the correct identification of *An. arabiensis, An. gambiae* and *An. coluzzii* mosquitoes constitutes an integral part of malaria vector control programmes, and insecticide resistance management.

A commonly used method for differential identification of *An. arabiensis*, *An. gambiae* and *An. coluzzii* involves a combination of protocols established by Scott *et al.* and Fanello *et al.*(Scott *et al.* 1993, Favia *et al.* 1997, Fanello *et al.* 2002). These methods are based on PCR-Restriction Fragment Length Polymorphism (PCR-RFLP) and make use of the presence of nucleotide substitutions within the 28S coding region, and part of the IGS region of rDNA (Fanello *et al*. 2002). More recently a Short Interspersed Nuclear Element (SINE) insertion (S200 X6.1) on the X chromosome of *An. gambiae* has been found to be fixed in all *An. coluzzii* and absent in *An. gambiae* and *An. arabiensis*. Additionally, a 26 bp deletion in the same region defines *An. arabiensis*, allowing for the development of a novel PCR diagnostic assay that differentiates the three species (Santolamazza *et al*.). Briefly, primers that flank the S200 X6.1 insertion were designed to amplify the genomic DNA isolated from *Anopheles gambiae s.s.* specimen. The PCR products are run on an agarose gel, and individuals (*An. coluzzii*) with the S200 X6.1 insertion show a single band at 479 bp while individuals (*An. gambiae*) with no S200 X6.1 insertion give a 249 bp band, and *An. arabiensis* 223 bp as a result of the deletion (Santolamazza *et al.* 2008). Either method for the identification of *An. arabiensis*, *An. gambiae* and *An. coluzzii* requires post-PCR analysis by gel electrophoresis, which is less sensitive than melt-curve analysis.

SYBR green DNA-based Real-Time PCR systems provide a good alternative to fluorescent probe-based Real-Time PCR techniques and are based on ability of SYBR green to produce a 100-fold increase in fluorescence when bound to double-stranded DNA. Even though SYBR green binds non-specifically to nucleic acids, the fluorescent signal produced when in complex with DNA is directly proportional to the length and amount of DNA copies synthesized during the reaction, making this technique very sensitive (Ho *et al*. 2010) and very precise when diagnostic primer sets are used.

The aim of this study is to demonstrate a time-efficient, highly sensitive and specific SYBR green-based real-time PCR diagnostic assay that differentiates between *An. arabiensis*, *An. coluzzii* and *An. gambiae*.

## MATERIALS AND METHODS

### Mosquito samples and DNA extraction

*Anopheles gambiae* mosquito samples were obtained from Vestergaard-NMIMR Vector Labs (VNVL) insectary. These consisted of more than 200 samples selected from the standard susceptible Kisumu strain originally from Kenya, and *An. gambiae* Tiassalé, a resistant strain from the village of Tiassalé in Côte d’Ivoire maintained in the VNVL insectary since 2010. Hundred and ninety *An. coluzzii* were obtained from Okyereko, a rice irrigation field in the Central region of Ghana (Okoye *et al.* 2005, Charlwood *et al*. 2012, Chabi *et al*. 2016). Hybrid *An. gambiae* / *An. coluzzii* (75 samples) were obtained from laboratory crossing of either *An. gambiae* Kisumu and *An. coluzzii* or *An. gambiae* Tiassalé and *An. coluzzii*. Fifty *An. arabiensis* samples were obtained from field collection is Kenya. The study was initiated at Noguchi Memorial Institute for Medical Research (NMIMR) in Ghana and the primer design, optimization, validation and high-throughput species identification at Liverpool School of Tropical Medicine (LSTM), UK.

Whole mosquito DNA extraction was performed using a simplified version of the protocol designed by Collins and colleagues (Collins *et al*. 1987). A single mosquito was homogenized in a 1.5 ml eppendorf tube containing 200 µl of CTAB buffer and incubated at 65°C in a water bath for 5 minutes. 200 μl of chloroform was added to the homogenate, mixed by inversion and centrifuged for 5 minutes at 12000 rpm 25°C. The supernatant was pipetted into new 1.5 ml eppendorf tubes. 200 μl of isopropyl alcohol was added, mixed by inversion and then centrifuged at 12000 rpm for 15 minutes. The supernatant was then discarded gently and the DNA pellet was thereafter purified with 70% ethanol, dried overnight, and reconstituted in 20 µl of DNAse free water.

### Mosquito species identification

*Anopheles arabiensis and An. gambiae* s.s. species were first determined using established protocols (Scott *et al*. 1993, Fanello *et al*. 2002). Conventional PCR assays were performed with a 1/40^th^dilution of the DNA obtained from a single mosquito. PCR products were run on 2% agarose gels, stained with ethidium bromide and then visualized using UV Transilluminator (BioDoc-It™ Imaging System).

SINE PCR was performed using primers designed by Santolamazza *et al*. (Santolamazza et al. 2008) for the identification of *An. arabiensis*, *An. coluzzii* and *An. gambiae* using the primers; F6.1a (TCGCCTTAGACCTTGCGTTA) and R6.1b (CGCTTCAAGAATTCGAGATAC). Total PCR reaction volume of 25 µl containing 4 µl of 1/40th dilution of genomic DNA (used as template), 1 µl each of 10 µM of both forward and reverse primers, 6.5 µl of nuclease-free water and 12.5 µl of GoTaq master mix (Promega, Madison, WI, USA). Reaction conditions used were 94°C for 10 minutes; followed by 35 cycles of 94°C for 30 seconds, 59°C for 30 seconds72°C for 1 minute; and a final extension at 72°C for 10 minutes. PCR products were run on 2% agarose gels, stained with ethidium bromide and thereafter visualized using UV Trans-illuminator.

### SYBR green-based Real-Time PCR for species identification

This designed melt curve method to distinguish between *An. arabiensis*, *An. gambiae* and *An. coluzzii* uses the diagnostic Short Interspersed Nuclear Element (SINE200). The original S200 X6.1 primers designed by (Santolamazza *et al.* 2008) were unsuitable for melt curve analysis since firstly they produce overlapping melt-peaks for *An. coluzzii* and *An. gambiae*/*An. coluzzii* hybrids and secondly they give indistinct melt peaks for *An. gambiae* and *An. arabiensis*. New primers were designed based on amplicon length and G/C content to produce distinct melt-curves for the three sibling species and the hybrids. A universal forward primer (SINE200Fa 5’-ATTGCTACCACCAAAATACATGAAA-3’) matching all three species is combined with *An. gambiae*/ *An. coluzzii* specific reverse (SINE200Rd 5’-GGGGGGGGGAATAATAAGGAACTGCATTTAAT-3’) and for *An. arabiensis* reverse (SINE200Re 5’- GGATGTCTAATAGTCTCAATAGATG-3’).

SINE200Rd, the reverse primer for *An. gambiae* and *An. coluzzii* has a G8 stretch added to the 5’ end of the primer (as underlined) to increase the amplicon melting temperature (Tm), preventing the melt profiles of *An. gambiae* and *An. arabiensis*to overlap. SINE200Rd incorporates the SINE200 transposon in *An. coluzzii* into the amplicon and gives a distinct high Tm. The specificity of the *An. arabiensis* primer SINE200Re is based on only two mismatches with *An. gambiae*. Both mismatches are conveniently on the 3′ end of the primer though a sufficiently high annealing temperature effectively prevents amplification of *An. gambiae.* The amplicon sizes are 60 bp, 103 bp and 333 bp for *An. arabiensis, An. gambiae* and *An. coluzzii* respectively.

A total of sixty samples including 10 *An. arabiensis*, 20 of *An. gambiae*, 14 *An. coluzzii*, and 15 hybrids randomly selected among the individuals characterized by SINE PCR, and one artificial hybrid (pooled DNA of *An. gambiae-An. coluzzii*) were analysed using the designed melt-curve protocol.

A total volume of 10 µl mixture was prepared by combining 3.75 µl of Nuclease-free PCR grade water, 5 µl Luna Universal qPCR Master Mix (New England Biolabs, Ipswich, Ma, USA), 0.25 µl of each (10 μM) SINE200Fa, SINE200Rd and SINE200Re primers, and 0.5 μl mosquito DNA template (Table 1).

**Table 1:** SYBR green Master Mix

|  | **1X (μl)** |
| --- | --- |
| **H<sub>2</sub>O** | 3.75 |
| **Luna Universal qPCR Master Mix** | 5 |
| **SINE200Fa (10 μM)** | 0.25 |
| **SINE200Rd (10 μM)** | 0.25 |
| **SINE200Re (10 μM)** | 0.25 |
| **Genomic DNA template** | 0.5 |
| **Total** | 10 |

The samples were run on AriaMx Real-time PCR system (Agilent, Santa Clara, Ca, USA) using the quantitative PCR DNA binding dye including standard melt curve program and 520 nm wavelength filter (FAM). Annealing temperature (Ta) and cycle number were optimised to eliminate non-target background melt-peaks. The optimized cycling conditions involved a denaturation of 95°C for 60 seconds followed by 33 cycles of 60°C for 20 seconds and 72°C for 10 seconds with a final dissociation step of 95°C for 60 seconds, 55°C for 30 seconds and a melt ramp up to 95°C with 0.5°C increments.

The robustness of the assay was tested using SYBR green-based alternative reagents and an alternative real-time PCR machine. Eight samples of each group (*An. arabiensis*, *An. gambiae*, *An. coluzzii*, and hybrid) were repeated with Luna Universal qPCR Master Mix and with Brilliant III Ultra-Fast SYBR Green Low ROX QPCR Master Mix (Agilent) on a Stratagene Mx3005P real-time PCR machine (Agilent). Thermal cycling conditions and PCR mix were identical to those used for AriaMX.

The species ID was automated using the following “nested if” statement in Microsoft Excel: IF(AND(A1=”N/A”, B1=“N/A”),“N/A”, IF(AND(A1>85,B1=“N/A”),“AC”, IF(AND(A1>85,B 1>74),“HY”, IF(AND(A1>74,B1=“N/A”),“AG”, IF(AND(A1>71,B1=“N/A”),“AA”))))).

With cell A1 containing the first melt peak and B1 (only present in case of a hybrid) the second melt peak. The peak temperature criteria are: >85°C = AC = *An. coluzzii*, between 85°C and 74°C = AG = *An. gambiae*, <74°C = AA = *An. arabiensis*, both >85°C and between 85°C and 74°C = HY = *An. gambiae /An. coluzzii* hybrid. This automation prevents the need of manually scoring species as is the case for gel-based species ID, thereby reducing time and scoring errors. A large number of samples (1075 mosquitoes) collected in western Kenya between 2011 and 2015 whose species ID was previously assigned using the 28S IGS gel-based method (Fanello et al. 2002) were re-examined using the melt-curve technique. A subset of 24 individuals identified as *An. gambiae* with the 28S method and *An. arabiensis* according to the melt-curves plus all 10 *An. arabiensis* re-scored as *An. gambiae* were repeated with the gel-based SINE200 method (Santolamazza et al. 2008).

## RESULTS

### Species identification

#### SINE PCR

The sixty samples selected were identified including 10 *An. arabiensis*, 20 *An. gambiae*, 14 *An. coluzzii* and 15 hybrids of both species and one pooled artificial hybrid. Gel electrophoresis analyses of SINE PCR products showed distinct diagnostic bands corresponding to each species. *Anopheles arabiensis*, *An. coluzzii* and *An. gambiae* showed specific band sizes of 315, 479 and 249 bp respectively, while the *An. gambiae*/*An. coluzzii* hybrids showed both band sizes. Figure 1 described the band sizes of some selected samples.

**Figure 1:**
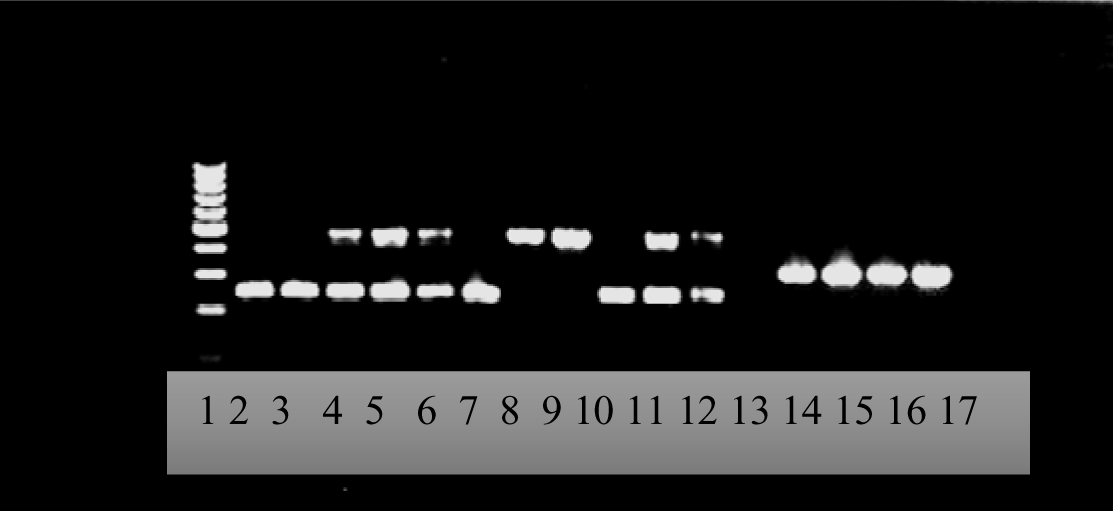
2.0% agarose gel for SINE PCR showing *An. arabiensis*, *An. gambiae* and *An. coluzzii Lane 1: molecular weight ladder (100 bp); Lane 2; 3 & 10: An. gambiae, Lane 4–6 & 11-12: hybrids An. coluzzii / An. gambiae; Lane 8 & 9: An. coluzzii, Line 14-17: An. arabiensis. DNA fragments of 315 bp for An. arabiensis, 249 bp (An. gambiae) and 479 bp (An. coluzzii). Hybrid showed both band sizes at 249 bp and 479 bp,*

#### SYBR green Real-Time PCR

The identification of all the species and hybrid was based on specific melting temperatures (Tm) from the dissociation curves (Figure 2). The Tm could slightly shift based of the DNA quality, extraction method or age and preservation of the mosquito samples. *Anopheles arabiensis* showed a single peak at an average temperature of 72°C*, An. gambiae* at 75°C; whilst *An. coluzzii* peaks at 86°C (Figure 2). The hybrid showed the expected 86°C melting peak for *An. coluzzii* and a slightly shifted 74°C peak for *An. gambiae*. The pooled *An. gambiae - An. coluzzii* DNA produced a melt-curve identical to hybrids. Results were consistent between the two Real-time PCR machines and between alternative SYBR green qPCR master mixes, except for a slight shift in species-specific melt peaks between the machines. This “machine effect” demonstrates the importance of using positive controls for the three species to calibrate the species-specific peak temperature intervals when using different real-time machines.

**Figure 2:**
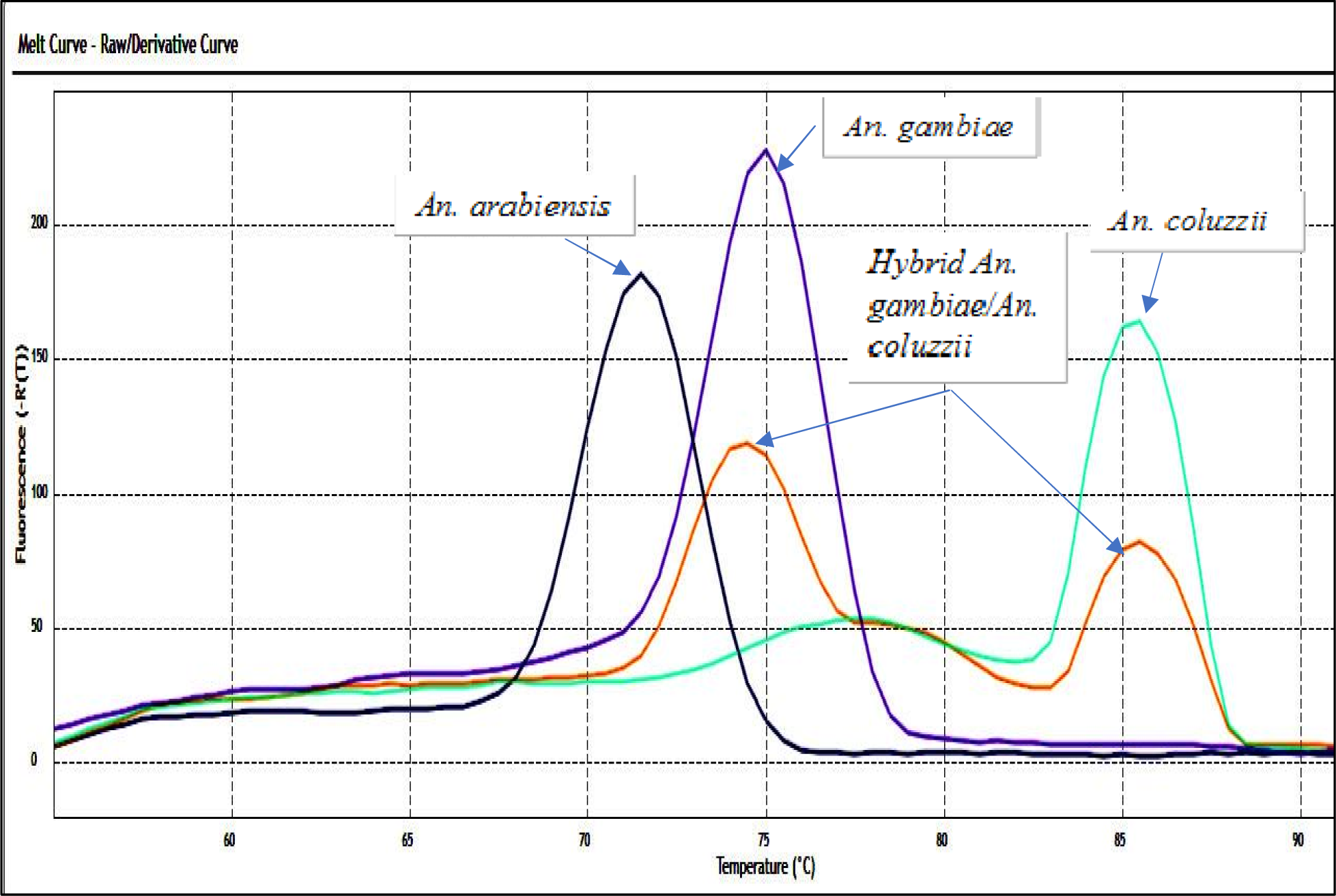
Dissociation curves of *all species* (*An. gambiae*, *An. coluzzii* and *An. arabiensis*) using SYBR green high throughput methods.

Re-evaluation of 1075 mosquitoes from western Kenya previously screened with 28S IGS revealed an error rate of 36% for the gel-based method. Out of 354 mosquitoes initially scored as *An. arabiensis*, 10 were identified as *An. gambiae* using the melt-curve technique, and 373 out of 721 assigned *An. gambiae* produced and *An. arabiensis*-specific melt peak. Verification of inconsistently scored mosquitoes using the SINE200 gel method confirmed the melt-curve species identification as the correct one.

## DISCUSSION

Rapid and reliable identification of species and sub-species of malaria vector populations is an important part of malaria vector control programmes. PCR-RFLP and SINE PCR methods have been developed for this purpose and have both shown to successfully identify *An. arabiensis*, *An. gambiae* and *An. coluzzii* out of the complex of eight species (Scott *et al*. 1993, Fanello *et al*. 2002, Santolamazza *et al*. 2008). However, both methods involve at least two PCR steps with gel staining that require precision and time to identify the species. In addition, inconsistent identification of *An. gambiae*, *An. coluzzii* and their hybrids has been reported by either PCR using form-specific primers or PCR-RFLP genotyping carried out in different laboratories (Santolamazza *et al*. 2011). Mis-identification of *An. gambiae* vs *An. arabiensis* based on gel-bands is even more likely since both the SINE200 and the 28S IGS techniques produce similar-sized amplicon sizes (223 vs 249 bp and 315 vs 390 bp respectively) which only separate when gels are run sufficiently long (the *An. gambiae*/*An. coluzzii* distinctive HhaI digest (Fanello *et al.* 2002) is usually not performed in Eastern Africa). The 36% error rate in western Kenyan samples clearly demonstrates the risk of mis-interpreting bands on gels, and consequently the value of the melt-curve approach described here. From Scott *et al*. 1993 to date, SINE PCR has shown greater reliability among all the protocols allowing the differentiation of *An. gambiae* and *An. coluzzii*. Moreover, each species identification protocol allows partial identification of the whole *An. gambiae* complex (Table 2). Though, the per-sample cost of SYBR green is 1.75 times higher than the reagents for gel-based species ID (£0.08 vs £0.14), but the benefits outweigh the extra expense given the faster and more reliable results.

**Table 2:**
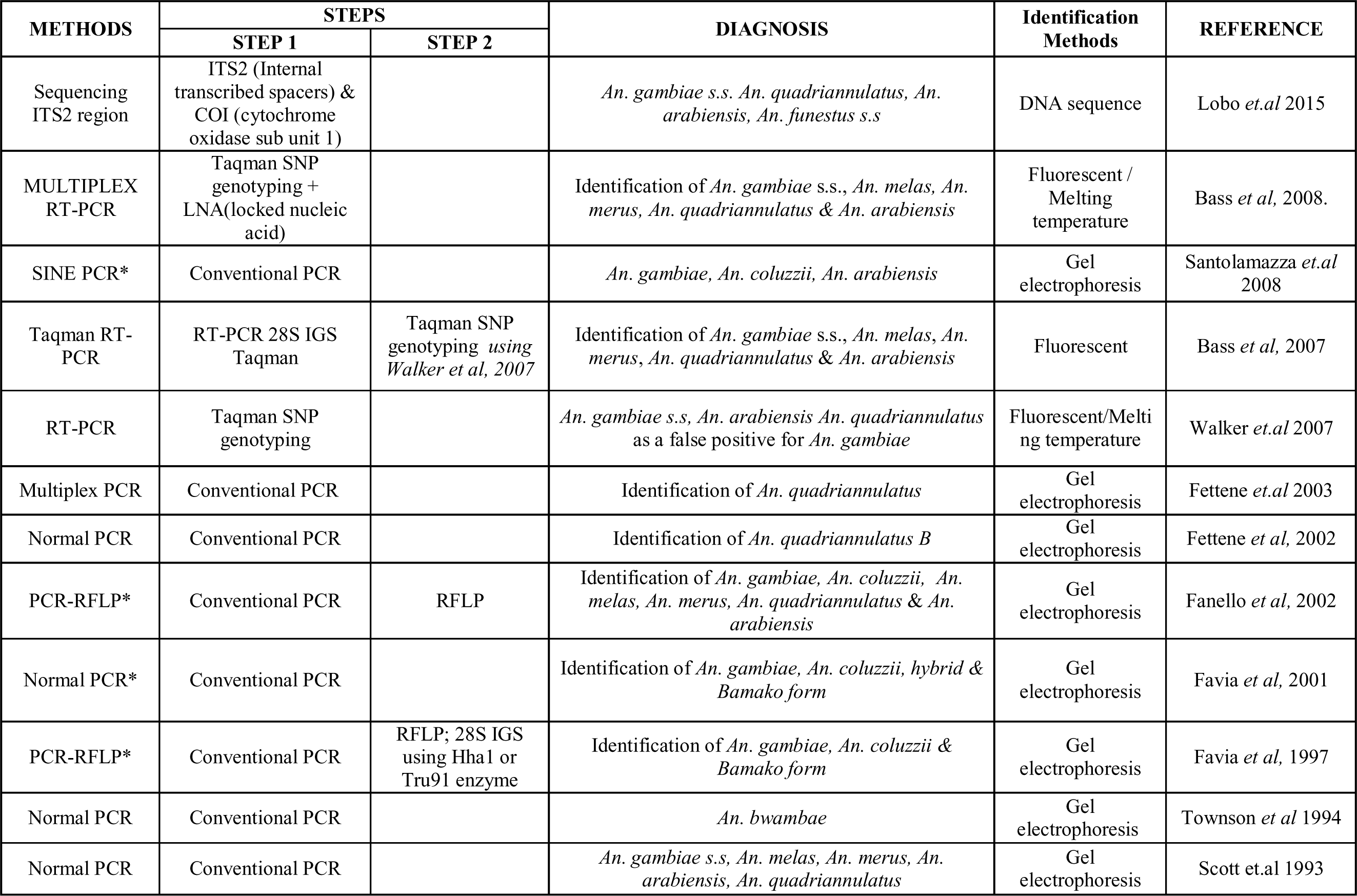
Summary of the different protocols for the identification of *An. gambiae* s.l complex [\**Represents the protocols enabling the identification of An. gambiae and An. coluzzii*]

| METHODS | STEPS |  | DIAGNOSIS | Identification Methods | REFERENCE |
| --- | --- | --- | --- | --- | --- |
|  | STEP 1 | STEP 2 |  |  |  |
| Sequencing ITS2 region | ITS2 (Internal transcribed spacers) & COI (cytochrome oxidase sub unit 1) |  | *An. gambiae* s.s. *An. quadriannulatus*, *An. arabiensis*, *An. funestus* s.s | DNA sequence | Lobo *et.al* 2015 |
| MULTIPLEX RT-PCR | Taqman SNP genotyping + LNA(locked nucleic acid) |  | Identification of *An. gambiae* s.s., *An. melas*, *An. merus*, *An. quadriannulatus* & *An. arabiensis* | Fluorescent / Melting temperature | Bass *et al*, 2008. |
| SINE PCR* | Conventional PCR |  | *An. gambiae*, *An. coluzzii*, *An. arabiensis* | Gel electrophoresis | Santolamazza *et.al* 2008 |
| Taqman RT-PCR | RT-PCR 28S IGS Taqman | Taqman SNP genotyping using Walker *et al*, 2007 | Identification of *An. gambiae* s.s., *An. melas*, *An. merus*, *An. quadriannulatus* & *An. arabiensis* | Fluorescent | Bass *et al*, 2007 |
| RT-PCR | Taqman SNP genotyping |  | *An. gambiae* s.s, *An. arabiensis* *An. quadriannulatus* as a false positive for *An. gambiae* | Fluorescent/Melting temperature | Walker *et.al* 2007 |
| Multiplex PCR | Conventional PCR |  | Identification of *An. quadriannulatus* | Gel electrophoresis | Fettene *et.al* 2003 |
| Normal PCR | Conventional PCR |  | Identification of *An. quadriannulatus B* | Gel electrophoresis | Fettene *et al*, 2002 |
| PCR-RFLP* | Conventional PCR | RFLP | Identification of *An. gambiae*, *An. coluzzii*, *An. melas*, *An. merus*, *An. quadriannulatus* & *An. arabiensis* | Gel electrophoresis | Fanello *et al*, 2002 |
| Normal PCR* | Conventional PCR |  | Identification of *An. gambiae*, *An. coluzzii*, hybrid & *Bamako form* | Gel electrophoresis | Favia *et al*, 2001 |
| PCR-RFLP* | Conventional PCR | RFLP; 28S IGS using Hha1 or Tru91 enzyme | Identification of *An. gambiae*, *An. coluzzii* & *Bamako form* | Gel electrophoresis | Favia *et al*, 1997 |
| Normal PCR | Conventional PCR |  | *An. bwambae* | Gel electrophoresis | Townson *et al* 1994 |
| Normal PCR | Conventional PCR |  | *An. gambiae* s.s, *An. melas*, *An. merus*, *An. arabiensis*, *An. quadriannulatus* | Gel electrophoresis | Scott *et.al* 1993 |

The aim of this study was to demonstrate a SYBR green, rapid, high throughput assay capable of identifying *An. arabiensis*, *An. gambiae*and *An. coluzzii* with high specificity and precision. The high throughput SYBR green rapid assay described here shows great specificity. Though the existing protocols have proven to be useful tools for species identification, this new high throughput assay does not require post PCR analyses such as restriction enzyme digestion and gel electrophoresis associated with PCR-RFLP and SINE PCR (Fanello *et al*. 20021, Santolamazza *et al*. 2008).

Thus, this new assay showed the first use of SYBR green designed set of primer sequences to distinguish *An. arabiensis*, *An. gambiae* and *An. coluzzii* by Real-Time melt-curve analysis. The newly developed method together with Taqman qPCR methods (Bass *et al.* 2008) will allow complete characterization of mosquito specimen into species using Real-Time PCR. Compared to the existing method, this assay is faster, and the closed-tube reaction (without gel electrophoresis) reduces the risk of post-PCR contamination. The assay designed for basic melt-curve analysis (rather than high-resolution melt-curve) allows the use on a wide range of real-time PCR machines.

## CONCLUSION

The SYBR green real time PCR techniques showed an additional option for the characterization of both sibling species and *An. arabiensis* while all the other real time PCR protocol of species differentiation could not allow the characterization of the sub-species such as *An. gambiae* and *An. coluzzii*. The assay designed in this study is a new tool to help researchers and particularly malaria vector control entities to identify clearly the subspecies of *An. gambiae* s.l within the complex in the context where each of the species has unique behaviour and impact on malaria.

## Acknowledgements

This work was financially supported by Vestergaard. We would like to acknowledge the contribution of Godwin Amlalo, Dorothy Obuobi and Rebecca Pwalia of Vestergaard-NMIMR Vector Labs for their support on part of the laboratory works. We thank David Weetman for his valuable comments on an early version of the manuscript.

## Author details

^1^Noguchi Memorial Institute for Medical Research, PO Box LG 581 Legon, University of Ghana, ACCRA Ghana, ^2^Liverpool School of Tropical Medicine, UK; ^3^Kemri-Wellcome Trust Research Programme, Kilifi, Kenya ^4^Vestergaard Frandsen East Africa, ABC Place 1st Floor, Waiyaki Way; City/Town, Nairobi, Kenya, ^5^Section Environmental Parasitology, Kobe-Tokiwa University, Japan; ^6^Vestergaard US regional office, 1020 19th Street NW |Suite 895 | Washington, DC 20036, USA.

## Authors’ contributions

JC, AVH and LKN designed, implemented the study, analysed and interpreted data. JC, AD and AVH drafted the manuscript. AVH, HN and DP improved the technique and designed the primers. DKDS, TS, SKD, MPH and HPJ revised the manuscript. AVH, LKN, HN, DP, AD and DO carried out all the laboratory experiments. All authors read and approved the final manuscript.

